# Rapid vegetation reorganization in Hengduan Mountains driven by high climatic oscillations

**DOI:** 10.64898/2026.06.23.734113

**Authors:** Mengna Liao, Pengyu Li, Ziru Hao, Xiao Zhang, Kai Cui, Yongbo Wang, Xingqi Liu, Enlou Zhang, Jian Ni, Kai Li

**Affiliations:** College of Life Sciences, Zhejiang Normal University, Jinhua, 321004, China; College of Resource Environment and Tourism, Capital Normal University, Beijing, 100048, China; Nanjing Institute of Geography and Limnology, Chinese Academy of Sciences, Nanjing, 211135, China

**Keywords:** Pollen analysis, Hengduan Mountains, rapid vegetation reorganizations, Marine Isotope Stage 2, AMOC, teleconnection

## Abstract

The accelerating global climate change has been triggering large-scale vegetation reorganizations, yet our understanding of how mountain ecosystems respond to rapid climatic oscillations is critically constrained. Here, we present a high-resolution palynological record from Erhai Lake, southwestern China, revealing eight episodes of rapid vegetation reorganizations in the Hengduan Mountains (HMs) over the past 35,200 years. These reorganizations closely tracked the rhythms of rapid climate oscillations, particularly during the Last Glacial Maximum and the Last Deglacial Period. We find that while the timing of rapid vegetation reorganizations were synchronous with Atlantic Meridional Overturning Circulation (AMOC) anomalies that modulated global climate variability, the magnitude of these reorganizations did not exhibit a linear correlation with AMOC strength; instead, they were governed by local heat and moisture availability mediated through teleconnections. This demonstrates a strong natural regulatory capacity of mountain vegetation in HMs, enabling resilience to intense climatic fluctuations. However, when using the Erhai record as a benchmark, we project that rapid reorganizations under the high-emission pathway (SSP585) will likely surpass the intensities observed during historical events. These findings reveal the high climatic sensitivity and strong natural regulatory capacity of mountain ecosystems, highlighting the critical necessity of climate mitigation actions and nature-based solutions to safeguard subalpine and alpine biodiversity against unprecedented future climate change.

## 1. Introduction

The accelerating rate of climate warming is raising the probability of abrupt ecosystem shifts worldwide^1, 2, 3^, resulting in large-scale reorganizations of terrestrial ecosystems^4, 5^. Montane ecosystems, characterized by high level of richness and endemicity of species, serve as indispensable refugia for lowland species and contribute disproportionately to the terrestrial biodiversity of Earth^6^. Assessing the extent to which mountain vegetation responds to rapid climate change will benefit our understandings on the vulnerability and sensitivity of mountain vegetation to global climate change and provide a crucial foundation for formulating targeted protection strategies.

The Hengduan Mountains (HMs), one of the world’s biodiversity hotspots^7^, harbor one of the planet’s richest temperate floras and support a diverse range of habitats and vegetation types^8^. Adjacent to the southeastern margin of the Tibetan Plateau, this region serves as a critical exchange point for global heat transfer and acts as a barrier that disrupts and modifies the flow of energy from low latitudes. Meanwhile, the north-south orientation of HMs creates a topographic corridor facilitating the penetration of the tropical moisture from the Indian Ocean into the plateau’s interior. These unique geographical and topographical conditions render the region closely linked to the global thermodynamic cycle through teleconnections and highly sensitive to global climate change^9, 10^. Thus, in the context of global warming, it is no surprise to observe significant warming trends in HMs over the past six decades^11, 12^. With climate warming, the treeline in HMs has migration upward by over 50 m in recent decades^13^, the vegetation coverage has been expanding notably^14^, and plant distribution has shifted widely^15^. This series of changes will inevitably promote alternations in plant community structure and composition, namely, vegetation reorganization. However, due to the lack of systematic investigations and long-term data on plant community, we rarely know the process and characteristics of vegetation reorganization that occurs in response to rapid climate change in this mountainous region. This will hinder the assessment of ecosystem stability and functioning, limit our understanding of ecological feedback mechanisms to climate warming, and impede the development of effective biodiversity conservation strategies for this critical biodiversity hotspot.

Geological records severed as natural experiments, enabling the studies of how mountain vegetation reorganizes, shifts its ranges, and adapts to rapid past climate changes, thereby offering a benchmark for predicting future vegetation changes ^16^. Numerous geological records demonstrate that HMs were influenced by a series of Northern Hemisphere-wide climatic events during the Last Glacial Maximum (LGM) within 27.0–18.0 calibrated thousand years before radiocarbon present (cal kyr BP, where 0 cal kyr BP is 1950 CE) and the Last Deglacial Period (LDP; 18.0–11.7 cal kyr BP). Key events include the Heinrich Stadial 1 (HS1, 17.5–15.0 cal kyr B), the Bølling–Allerød warming period (B/A; 15.0–13.0 cal kyr BP), and the Younger Dryas (YD; 13.0-11.7 cal kyr BP) ^17, 18, 19, 20^. These climate events act as a natural window that allow us to prob deeply into how vegetation responses to rapid climatic shifts. Based on palaeoecological records (primarily subfossil pollen from lake sediments), we know that vegetation in HMs underwent widespread and noticeable changes in response to these abrupt climate shifts ^17, 19, 20, 21, 22^. These studies reveal some common patterns in regional vegetation responses to rapid climate events, such as vertical shifts in vegetation zones and treelines, as well as changes in the openness of alpine vegetation that correspond to rapid warming or cooling. However, it remains uncertain how vegetation in response to rapid climatic shifts—whether through gradual reorganization (e.g., spanning more than a century) or rapid reorganization (e.g., occurring over a few decades). This uncertainty partially arises from the scarcity of high-resolution paleoecological records in the region that are capable of capturing details of vegetation zone shifts.

Here, we present a high-temporal-resolution (80 years in average) palynological record (EH22) from the central part of Erhai Lake in the southern HMs (Fig. 1, Fig. S1). This core is supported by a well-constrained chronology spanning the past 35,200 years (Fig. S2). With this new record, we identify multiple episodes of rapid vegetation reorganization during the LGM and LPD. By diagnosing the background climatic conditions associated with these vegetation changes, we demonstrate that these rapid reorganizations were significantly modulated by global climate teleconnections driven by large-scale latitudinal thermal exchanges. Furthermore, we assess the possibility of future rapid vegetation reorganization under various Shared Socioeconomic Pathway (SSP)-based scenarios, using the reconstructed patterns of past rapid vegetation organizations as a benchmark. Collectively, these analyses reveal high climatic sensitivity of HMs vegetation across both paleo and future contexts, offering novel insights for developing adaptive conservation strategies in mountain ecosystems.

**Figure 1.**
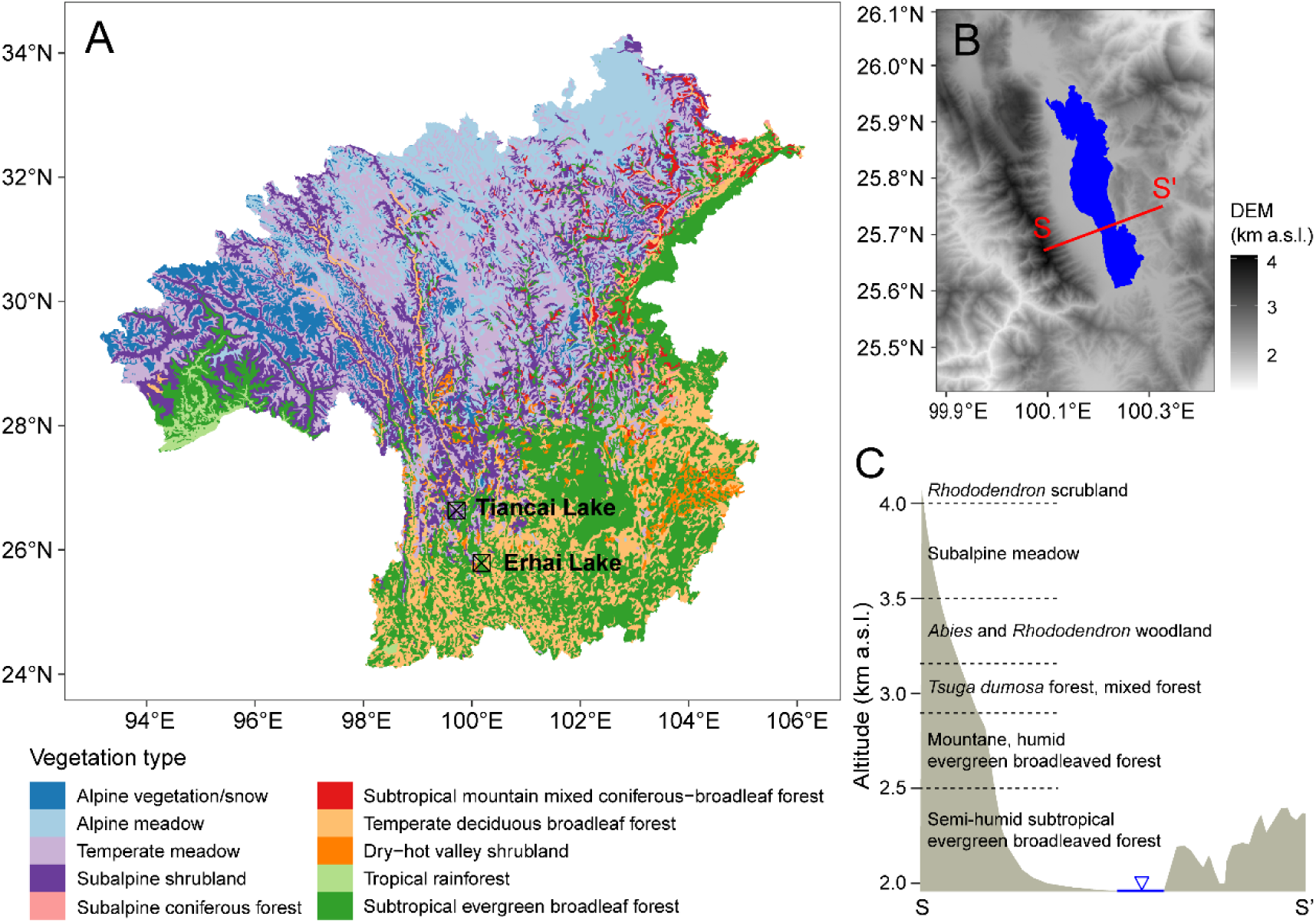
Geographic context and vegetation patterns in the study region. A, Location of the Erhai Lake, with Tiancai Lake in the north (referred summer temperature record in main text^9^. The background map illustrates the zonal vegetation patterns across the Hengduan Mountains^23^. B, The topographic features of the Erhai Lake Basin. C, Vertical vegetation zonation in the Erhai Lake Basin along the section S-S’ depicted in B.

## 2. Results and Discussion

### 2.1 Vegetation dynamics over the past 35,200 years

Pollen assemblage of core EH22 mainly comprises *Pinus*, *Quercus*, *Tsuga*, *Picea*, and *Abie*s (Fig. 2, Fig. S3), which are constructive taxa in pine forest, broadleaved forest, hemlock forest, and cold coniferous forest, respectively^23^. Despite their mixed growth to a certain extent, these forests primarily occupy different vertical vegetation zones^24, 25^. Nowadays, the broadleaved forest mainly distributes below 2900 m above sea level (a.s.l.), the hemlock forest ranges between 2900–3200 m a.s.l., and cold coniferous forest is prevailing from 3200 to 3500 m a.s.l. in HMs (Fig. 1). Therefore, the dynamics in pollen spectra from core EH22 can be interpreted as vertical migration of vegetation zones and the reconfiguration of plant communities in the southern HMs over the past 35,200 years.

**Figure 2.**
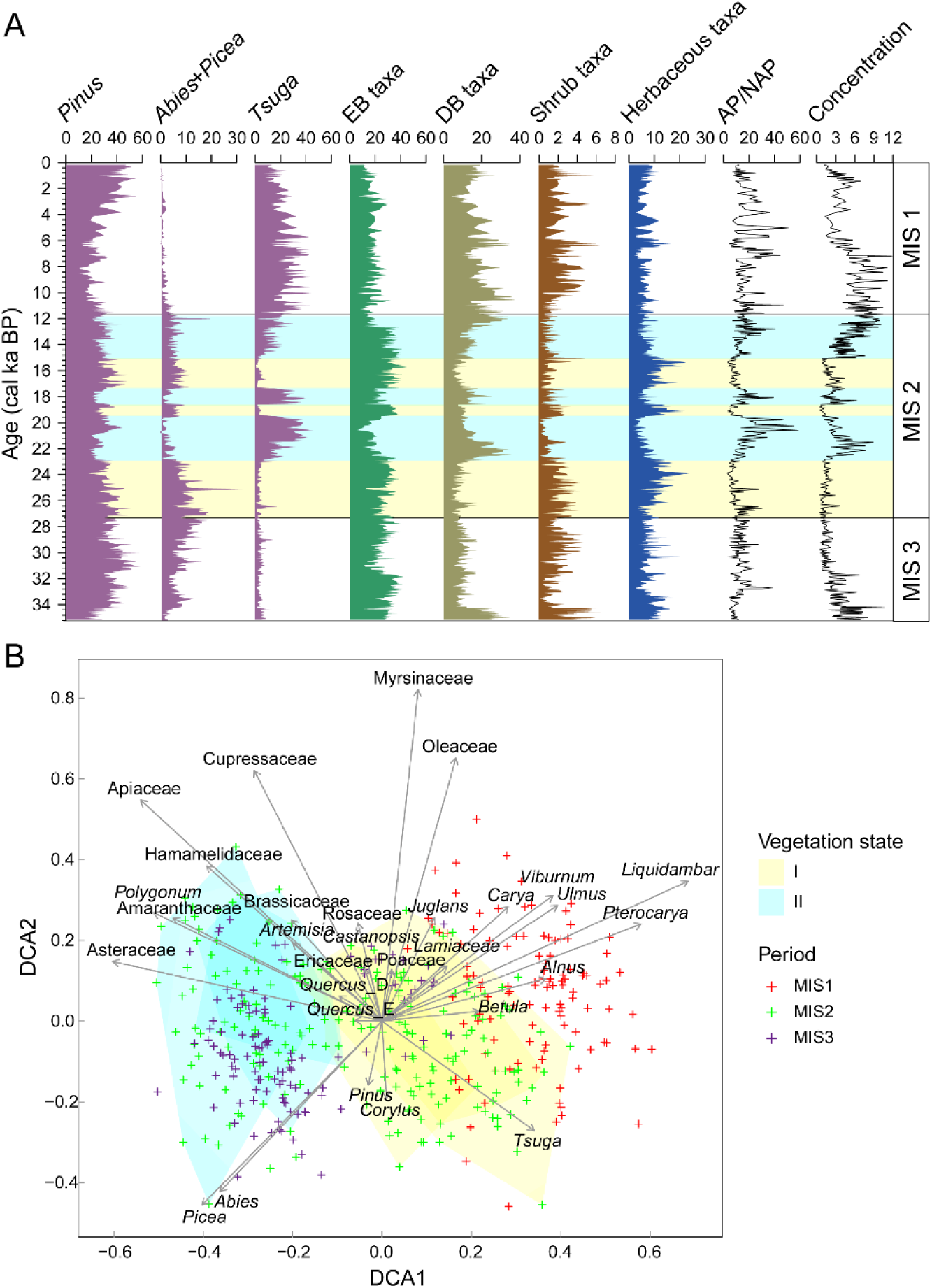
Pollen assemblage change over the past 35 cal kyr BP. **A,** Temporal trends in pollen group percentages, arboreal/non-arboreal pollen ratio (AP/NAP), and pollen concentration (con.). Evergreen broadleaf (EB) taxa: evergreen *Quercus*, Euphorbiaceae, *Ilex*, *Castanopsis*, Sapindaceae, Symplocaceae, Bignoniaceae, Magnoliaceae, *Mallotus*. Deciduous broadleaved (DB) taxa: *Betula*, *Juglans*, *Carya*, *Ulmus*, *Alnus*, deciduous *Quercus*, Hamamelidaceae, Tiliaceae, *Corylus*, *Carpinus*, Anacardiaceae, *Liquidambar*, Sterculiaceae, *Fagus*, *Salix*, Bombacaceae, Moraceae, *Celtis*, *Pterocarya*. Shrub taxa: Myrsinaceae, Celastraceae, Araliaceae, Thymelaeaceae, Rosaceae, Ericaceae, Campanulaceae, Oleaceae, *Lonicera*, *Viburnum*, Nitrariaceae, Rhamnaceae, Vitaceae, Myrtaceae, *Alangium chinense*, *Phyllanthus*. Herbaceous taxa: Poaceae, *Polygonum*, *Artemisia*, Asteraceae, Rutaceae, Acanthaceae, Elaeagnceae, Caryophyllaceae, Amaranthaceae, Brassicaceae, Lamiaceae, Liliaceae, Polygalaceae, Solanaceae, Apiaceae, *Mimosa*, Fabaceae, Onagraceae, Boraginaceae, Scrophulariaceae, Plantaginaceae, Verbenaceae, Rubiaceae, Gentianaceae, Balsaminaceae, Geraniaceae, Ephedraceae. **B,** Detrended correspondence analysis (DCA) of pollen assemblage, excluding taxa with mean abundance < 0.1%.

During the late Marine Isotope Stage (MIS) 3 (35.2–27.5 cal kyr BP), a moderate climate has been recorded in the southwestern China^26, 27, 28^, with mean annual temperatures (MAT) and precipitations (MAP) of 12℃ and 1150 mm, respectively, in the southern HMs. Accordingly, the vegetation remained stable in the study area, comprising mainly pine forest and evergreen oaks forest (Fig. 2A). The increasing trend in the abundance of *Abies*, *Picea* and shrub taxa indicates an expansion of cold coniferous forest and alpine shrubland, which is likely attributed to the decreasing temperature and precipitation in this period^26^. During the LGM, the atmospheric CO2 concentrations were about 100–120 ppm lower than pre-industrial Holocene levels^29^. MAT and MAP in the southern HMs decreased to 9℃ and 950 mm, respectively^26^. Meanwhile, the India Summer Monsoon (ISM) weakened obviously within 27–24 cal kyr BP but revived after that^30^. The pollen assemblages indicate an expansion of herbaceous vegetation during the early LGM, followed by rapid expansion or contraction of hemlock forest and deciduous broadleaved forest since 23 cal kyr BP (Fig. 2A). The expansion of cold coniferous forest and subalpine shrubland was likely restricted during this period. Both the atmospheric CO2 concentrations and global temperature rose to near pre-industrial levels during the LDP^31^. In HMs, this climatic transition was characterized by pronounced seasonal and annual post-glacial warmings^9, 26, 27^ and the intensification of the ISM^30^, which facilitated the expansion of evergreen broadleaved forest (Fig. 2A). The Holocene climate was characterized by a relatively high stability, with MAT around 17℃ and MAP exceeding 1300 mm in the southern HMs^32^. These climatic conditions facilitated long-term, progressive community reconfiguration, ultimately leading to the dominance of hemlock and deciduous broadleaved forests within Erhai catchment (Fig. 2A) and through the southern HMs^19, 20^.

### 2.2 Rapid vegetation shifts during the MIS 2

Our pollen record discloses frequent and rapid vegetation reorganizations during the MIS 2 (27.5–11.7 cal kyr BP), which contrasts with the stable vegetation state observed during the MIS 1 (11.7 cal kyr BP to the present) and the late MIS 3 (Fig. 2A). The climate during the MIS 2 was characterized by pronounced climatic variability, marked by the abrupt recurrence of millennial-scale cold-dry stadials alternating with warm-humid interstadials^33^. Coincidentally, in tandem with the rhythm of rapid climate change during this period, vegetation in the southern HMs underwent dramatic shifts between two distinct states (Fig. 2B). These shifts involved changes in plant populations and the reconstruction of plant communities over just a few decades (ranging from 20 to 50 years, determined by the time interval between the adjacent sediment samples).

The difference between these two vegetation states can be depicted by the Detrended Correspondence Analysis (DCA), an ordination method employed to uncover trends of variation in the composition of assemblages under environmental pressure gradient. Vegetation State I is characterized by relatively high proportions of *Tsuga*, *Alnus*, *Betula*, and *Carya* (Fig. 2B), along with a high ratio of arboreal to nonarboreal pollen (AP/NAP) and a high pollen concentration (Fig. 2A). However, in State II, these taxa are largely replaced by *Abies*, *Picea*, evergreen *Quercus* (Fig. 2B), and various herbaceous taxa, accompanied by a decline in the AP/NAP ratio and a reduction in pollen concentration (Fig. 2A). Positive scores along the first axis of DCA (DCA1) are associated with deciduous taxa and *Tsuga* (Fig. 2B), which represent non-alpine vegetation types. In contrast, negative scores along DCA1 are primarily driven by *Abies*, *Picea*, shrub and herbaceous taxa (Fig. 2B), largely indicative of alpine vegetation types. Based on the rate of change (RoC) in the DCA1 sequence, eight rapid shifts in pollen assemblage have been identified (Methods), with the majority of these shifts occurring during the MIS 2 (Fig. 3A). The rapid replacement or rebound dynamics between alpine and non-alpine taxa during the MIS 2 suggest that rapid vegetation reorganizations within the Erhai basin are closely related to the context of high climatic variability. This finding suggests that vegetation in HMs show high variability during cold intervals and low variability within the warm post-glacial period, a pattern consistent with the pronounced climate variability observed during the LGM and the subsequent global decline in climate variability from the LGM to the Holocene^34, 35, 36^.

**Figure 3.**
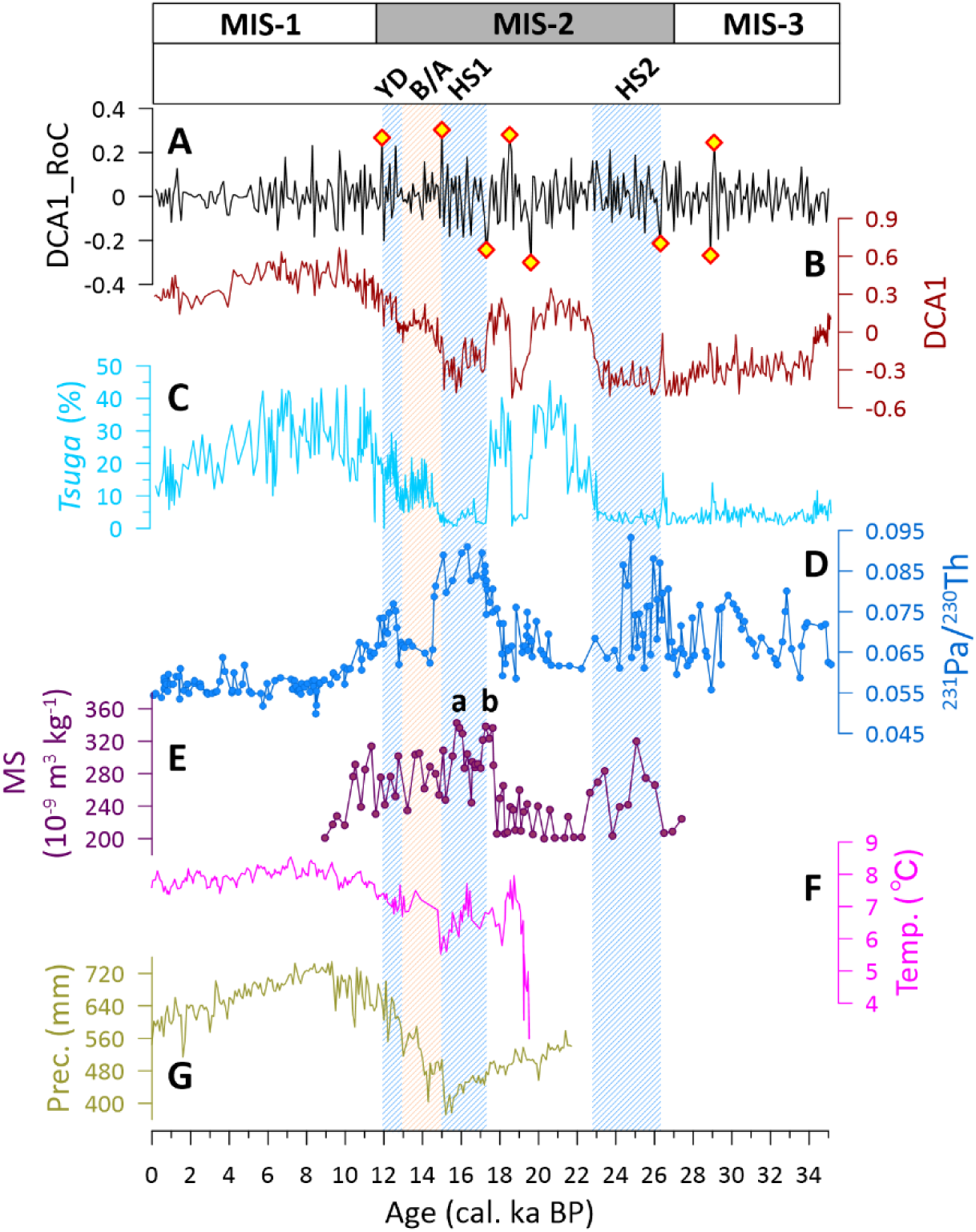
Rapid vegetation shifts and teleconnections with global climate. **A,** rate-of-change (RoC) of the 100-year interpolated DCA1 sequence and eight rapid vegetation shifts; **B,** DCA1 scores of pollen assemblages; **C,** *Tsuga* pollen abundance; **D,** Bermuda Rise ^231^Pa/^230^Th data^37^; **E,** Tore Seamount magnetic susceptibility (MS) record^38, 39^; **F,** Chironomid-inferred mean July temperature record from Tiancai Lake^9^; **G,** Simulations of precipitation during warmest quarter in the Erhai region (25.42–26.42°N, 99.88–100.43°E)^40^.

Although thermal deficiency exerts climate pressure globally during the MIS 2, vegetation responses show considerably spatial variability within HMs, with distinct ecological reactions to climate fluctuations at different locations^18, 19, 20, 41^. To date, no other published pollen records from HMs have been demonstrated such pronounced, rapid shifts in pollen assemblages during the MIS 2 as those documented in our record. Apart from climatic pressure, the availability of habitats severely restricts the impact of climate on alpine vegetation at high altitudes^42^, whiles in low-altitude areas, climate variability is significantly constrained by terrain features^14^. This partially explains why abrupt climate changes and rapid vegetation shifts do not align temporally in the high- and low-altitude regions of HMs. West of Erhai Lake rises the imposing Cangshan Mountain, reaching an altitude of approximately 4,100 m a.s.l. (Fig. 1B). It gradually extends northward to connect with the Tibetan Plateau, forming a pronounced ecological transition between the Yunnan lowlands and high-altitude terrain of HMs. This ecological transition zone fosters complex vegetation diversity by maintaining intact migration corridors and providing sufficient elevational refugia^24, 25^. These features enable rapid altitudinal migrations in plant populations, as well as community reorganization in response to rapid climate change^43^. In our record, the rapid alternation between two distinct vegetation stages is most prominently reflected in the relative abundance of *Tsuga* (Fig. 2A, Fig. S3). Consequently, our pollen data provide robust evidence suggesting that the hemlock forest in HMs demonstrates the highest sensitivity to rapid climatic fluctuations.

### 2.3 Potential mechanisms driving rapid vegetation reorganization

Rapid ecological changes are common phenomena in natural ecosystems which arise from a variety of circumstances, including rapid changes in external drivers (e.g., climate pressure, or resource constraints), nonlinear responses to gradual changes in drivers, and interactions among multiple drivers and disturbances^1^. It is evident that the timing of the rapid vegetation organizations, as indicated by the RoC of the DCA1 sequence, in HMs coincides with the onset and/or termination of North Atlantic climate events including HS2 (26.5–24.3 cal kyr BP), HS1, B/A, and YD (Fig. 3A). Notably, both the DCA1 sequence and the percentage abundance of *Tsuga* show fluctuations that closely track the intensity variations of the Atlantic Meridional Overturning Circulation (AMOC)^37, 44^ (Fig. 3D).

The AMOC that modulates oceanic poleward heat flux was less stable during the LGM than during the Holocene^37^. A weakened AMOC would reduce northward heat transport, triggering global-scale alternations in temperature and precipitation regimes and consequently leading to heat accumulation in low latitudes and a weakening ISM^9, 45, 46^. Accordingly, in our record, the cool- and moist-affinity taxa, e.g., *Tsuga*^47^, showed a synchrony decline concurrent with the slowdown of the AMOC, while the warm- and drought-tolerant taxa, e.g., sclerophyllous *Quercus*, increased (Fig. 1, Fig. S3). A similar pattern has been recently documented in northern Amazon forests during the HS1, where heat accumulation due to the slowdown of the AMOC was evidenced by a decline in cold- and moist-affinity vegetation elements, coupled with an increase in seasonal tropical vegetation^48^. Additionally, the rapid vegetation reorganizations within the HS1 corresponds well to a two depositional phases of ice-rafted detritus centering at 16.0 (HS1a) and 17.5 (HS1b) cal kyr BP^39^ (Fig. 3E). Conversely, the resumption of the AMOC promotes poleward heat transport and induces a northward shift of the monsoonal rain belt, which likely trigged the rapid expansion of hemlock forest in HMs during the MIS 2 and the Holocene (Fig. 3C). Interestingly, our pollen record identifies two rapid vegetation reorganizations that occurred between 20 and 18 cal kyr BP (Fig. 2A); however, this interval lacks well-documented global climate events to explained such rapid reorganization. Nevertheless, ^231^Pa/^230^Th data indicate a slowdown of the AMOC during this period, albeit with a magnitude of change less pronounced than that observed during HS1 (Fig. 3D). This suggests that large-scale oceanic-atmospheric interactions underwent alternations to a certain extend. A proxy-based temperature reconstruction from Tiancai Lake (Fig. 1A) reveals a marked rise in air temperature from between 19.5 and 18.5 cal kyr BP, likely surpassing temperatures recorded during the HS1 (Fig. 3F)^9^. These findings imply that the rapid vegetation shifts observed in HMs were directly induced by local availability of both heat and moisture, which in turn were regulated by global oceanic-atmospheric interactions.

The rapid vegetation reorganizations recorded in our pollen data were characterized by either sharp declines or rapid rebounds in the abundance of *Tsuga* (Fig S. 3A–3C). *Tsuga* is a taxonomic group that thrives in shaded and humid environments and is vulnerable to prolonged droughts^47^. Hemlock forests, which are predominantly composed of this genus, are commonly observed within cloud belt zones of HMs, where persistent fog and cloud cover help maintain soil moisture and reduce evapotranspiration. Therefore, the rapid reorganizations in HMs were likely regulated by relative humidity. Relative humidity is defined as the ratio of the actual amount of water vapor in the air to saturation capacity at a given temperature, primarily governed by air temperature. Although HMs experienced relatively low precipitation levels during the LGM (except for the interval of 19.6–18.5 cal kyr BP), the cold air temperature reduced saturation vapor pressure and likely maintained high relative humidity. Such climatic conditions likely resulted in an increase in relative humidity, consequently favoring the growth and reproduction of *Tsuga* species and the expansion of hemlock forests. In contrast, during HS1 and the interval from 19.6 to 18.5 cal kyr BP, warming temperatures coupled with reduced precipitation created climatic conditions characterized by persistent low relative humidity, which inevitably constrained the survival of *Tsuga* species.

### 2.4 Future vegetation prediction in Hengduan Mountains

Modern climate warming is amplifying global climatic variability, including the likelihood of abrupt climate changes^49^. This raises a critical question: under projected warming scenarios, will HMs experience rapid vegetation reorganization as intense as those observed during the MIS 2? Based on the topological structure parameters (including degree, betweenness, closeness, and eigenvector centralities) calculated for the major terrestrial pollen types (Methods), *Tsuga* was diagnosed as the pivotal node within the pollen assemblage network (Fig. S4). This indicates that changes in the taxon can reflect overall shifts in vegetation structure and composition. Therefore, to address the above question, we simulated the potential distribution of *Tsuga* across the entire HMs using Species Distribution Models (SDMs) for four future periods (2021–2040, 2041–2060, 2061–2080, and 2081–2100 CE) under four climate scenarios (SSP126, SSP245, SSP370, SSP585) (Methods).

The SDMs show that, by the end of this century, the habitats that are mediumly to highly suitable for *Tsuga* colonization within the Erhai catchment are projected to consistently increase, with a probable rapid expansion phase occurring during 2021–2040 CE (Fig. 4, Fig. S5). Notably, under the SSP585 scenario, the projected change in the proportion of medium to high habitats suitability between 2021–2040 and 1970–2000 CE (a difference of 0.26) is greater than the average change (0.2) that occurred during the vegetation transition from State I to State II during 22.0–12.6 cal kyr BP (Fig. 4, Fig. S6). These findings indicate that future climate change will drive rapid shifts in *Tsuga* distribution within the Erhai region. Such distributional changes are likely to disrupt existing ecological balances, reshuffling community compositions and potentially resulting in the emergence of novel communities^50, 51^. Consequently, the Erhai region may experience rapid vegetation reorganization in the near future. When examining the entire HMs, it is evident that suitable habitats for *Tsuga* are projected to change widespread, especially from the central to northern HMs, with the most pronounced changes occurring under the SSP585 scenario (Fig. 4, Fig. S6). This indicates that HMs may experience widespread rapid vegetation reorganization in the near future, despite exhibiting pronounced spatial heterogeneity.

**Figure 4.**
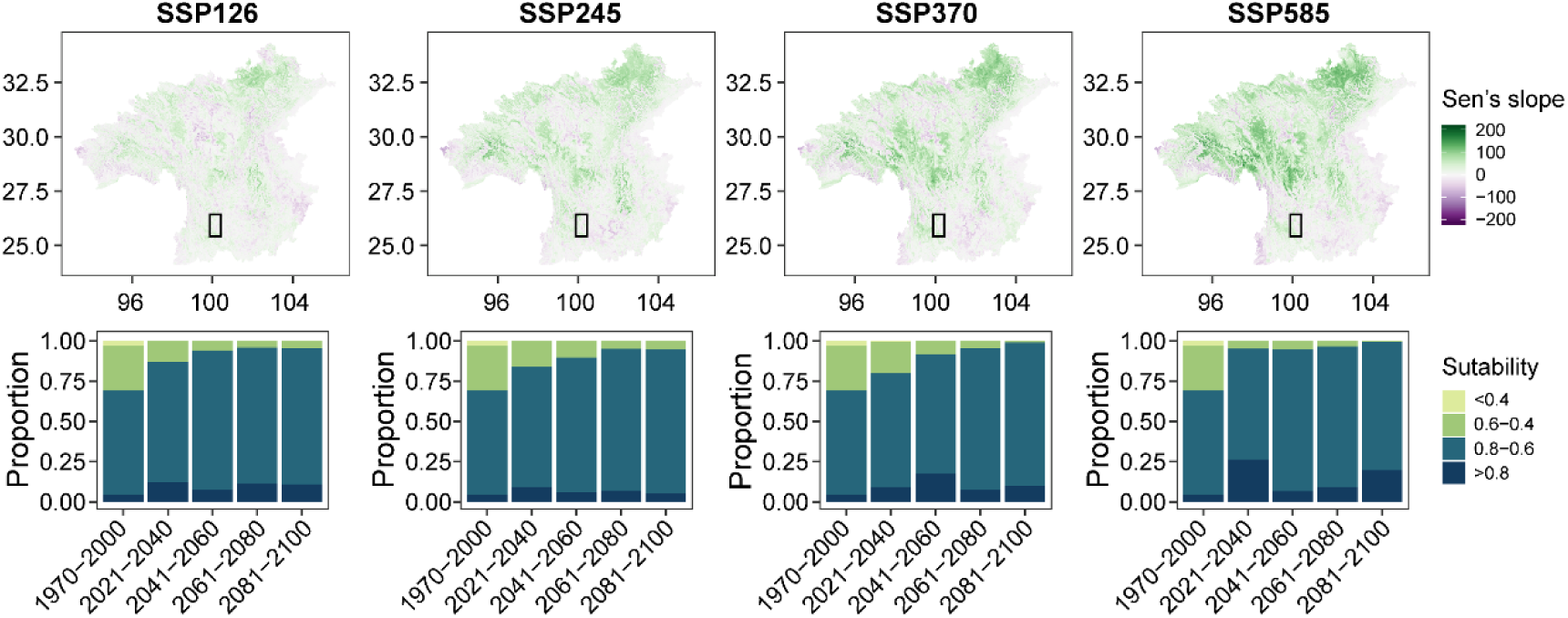
Projected trends in *Tsuga* habitat suitability across four consecutive periods (2021–2040, 2041–2060, 2061–2080, 2081–2100 CE). **Upper panel,** Sen’s slope estimators of *Tsuga* habitat area changes in the Hengduan Mountains under four Shared Socioeconomic Pathway (SSP) scenarios (SSP126, SSP245, SSP370, SSP585). **Lower panel,** Proportions of *Tsuga* habitat suitability levels in Erhai region, corresponding sequentially to the four climate scenarios depicted in Upper panel. The spatial extend of Erhai region is delineated by the rectangle in Upper panel.

In conclusion, this study demonstrates that montane vegetation in HMs exhibits exceptionally high climatic sensitivity, as evidenced by rapid vegetation reorganizations during the MIS 2. The occurrence of the rapid vegetation organizations is closely linked to the North Atlantic climate events, including HS2, HS1, B/A, and YD, indicating that these reorganizations were primarily triggered by climatic conditions characterized by high variability. Changes in the strength of the AMOC was identified as the core mechanism driving past rapid vegetation organizations in HMs. However, the magnitude of these reorganizations was not linearly correlated with AMOC strength. Instead, it was directly modulated by local heat and moisture availability through teleconnections associated with large-scale oceanic-atmospheric interactions. Our pollen record recognizes *Tsuga* as the indicator taxon that reflecting rapid vegetation reorganization. Under future scenarios, projections of *Tsuga*’s potential distribution indicate that the Erhai region, along with extensive areas of HMs, will undergo rapid vegetation reorganizations, with intensities under high-emission pathway (i.e., SSP585) likely exceeding those observed during historical events. Our study demonstrates that the mountain vegetation in HMs possess strong natural regulatory capacity. This underscores the need to prioritize nature-based solutions for conserving subalpine and alpine ecosystems in this region. Furthermore, given the critical role of hemlock (*Tsuga* spp.) forests as ecological nodes within the local vegetation system and their acute sensitivity to climatic variability, enhanced conservation efforts are needed to protect these forests. Although controversies exist regarding the recent collapse of the AMOC ^52, 53^, projections indicate that a continued slowdown of this oceanic circulation system will likely prevail in the coming decades ^54^. This situation warrants particular attention, as our pollen record reveals that even without a complete AMOC shutdown, the nonlinear response of local climates to moderate AMOC weakening could still trigger drastic vegetation reorganization in HMs.

## 3. Materials and Methods

### 3.1 Study site

Erhai Lake (25°36’–25°58’N, 100°06’–100°18’E, 1922 m a.s.l.) is located in the southwestern HMs on the southeastern Tibetan Plateau (Fig. 1A). The lake basin has an elongated shape and the major axis is oriented in a north-south direction. Erhai Lake covers an area of 249 km^2^, with a maximum water depth of 21 m and an average depth of 10 m^55^. Climate in this area is dominated by plateau subtropical monsoon climate, characterized by cold and dry winters and warm and humid summers. Observations from Dali Meteorological Station (25°42’N, 100°11’E, 1990.5 m a.s.l.) situated within the catchment yield a mean annual temperature of 15.1°C, a mean annual precipitation of 1055 mm, and around 85% precipitation falling between April and October (China Meteorological Data Service Centre, http://www.nmic.cn/).

The lake is surrounded by mountains, and the natural vegetation within its catchment area exhibits a distinct vertical distribution (Fig. 1C). The zonal vegetation is dominated by semi-humid subtropical evergreen broadleaved forest, primarily distributed at altitudes between 1,700 of 2,500 m a.s.l.. The arboreal layer of these forests is mainly composed of drought-adapted species from the genus *Cyclobalanopsis*, *Castanopsis*, and *Lithocarpus*. As altitude increases above 2500 m a.s.l., the subtropical evergreen broadleaved forests are gradually replaced by montane, humid evergreen broadleaved forests, which are predominantly dominated by *Lithocarpus* species. Species from *Castanopsis*, *Cyclobalanopsis*, *Schima*, and *Skimmia* are also commonly present in this zone. Above 2800 m a.s.l., with increasing moisture availability, the montane, humid evergreen broadleaved forests transition to hemlock (*Tsuga Dumosa*) forests and evergreen conifer-broadleaf mixed forests. Beyond 3,200 m a.s.l., *Abies* and *Rhododendron* woodlands, subalpine meadows, and *Rhododendron* shrubland successively emerge, forming a well-defined vertical spectrum of vegetation zones.

### 3.2 Field sampling, sediment lithology, and dating

A sedimentary core of EH22 (100.1795°E, 25.7892°N, 1542 cm in length) was retrieved from Erhai Lake on April 2022, at a water depth of 20 m using a UWITEC sample system. The lithology of the sediment profile is highly homogeneous upon visual inspection, primarily comprising greyish clayey silt with no discernible bedding structures or evidence of depositional hiatus (Fig. S1). A total of 14 samples from different depths were extracted for AMS ^14^C measurement in the Beta Analytic (SGS Beta), including 13 organic-sediment samples and one plant remain (Table S1). A carbon reservoir effect of 1160 years was estimated based on results of organic sediment and plant remain from the same depth of 1230 cm. The age-depth model of EH22 was created by using the R package *rbacon*^56^, with the IntCal20 radiocarbon calibration curve^57^. According to the age model, the basal sediment was deposited between 34.62–35.87 cal kyr BP with a median age of 35.2 cal kyr BP. The sedimentary rate was stable, averaging at 22.8 yr cm^−1^ (Fig. S2).

### 3.3 Pollen analysis

A total of 430 samples for pollen analysis were treated using the heavy liquid floatation method^58^ involving four main steps: removing carbonate with 10% HCl, removing humic substances with 10% KOH in a water bath of 80°C, performing heavy liquid flotation with a zinc bromide solution (ZnBr2 at approximately 2.2 g ml^−1^), and removing cellulose with acetolysis. To calculate the concentrations of pollen, tablets containing a known quantity of *Lycopodium* spores (10,315 grains per tablet) were added to each sample prior to the treatments. Pollen counts and identification were conducted using a ZEISS Lab.A1 microscope. At least 300 terrestrial pollen grains were count for each sample. Pollen identification was referred to the Chinese pollen books^59, 60^. Pollen percentages were calculated based on the total number of pollen grains from terrestrial pollen taxa and used to construct a pollen diagram using the Tilia software^61^. It is worth noticing that, to enhance the detection precision of rapid vegetation shifts, we increased sample density in stratigraphic layers showing significant changes in pollen assemblages. According to the established chronological framework (Fig. S2), all identified rapid vegetation shifts (Fig. 3A) occurred within 20–50 years, except for the event at 19.6 cal kyr BP, which spanned approximately 70 years.

### 3.4 Ordination analysis and abrupt change detecting

To determine the presence and timing of rapid changes in pollen data we firstly conduct Detrended Correspondence Analysis (DCA) using R package *vegan*^62^. Pollen percentage data were square-root transformed before conducting DCA to stabilize their variances. We selected the scores of DCA axis 1 (DCA1) to estimate the RoC in pollen assemblages, as this axis captures the greatest proportion of compositional variations in pollen data. The time sequence of DCA1 was interpolated at 100-year intervals. We calculated the RoC using the interpolated DCA1 sequence by determining the difference between consecutive data points. Rapid changes were finally identified by RoC values exceeding the 99^th^ percentile or falling below the 1^st^ percental of the interpolated DCA1 sequence.

### 3.5 Co-occurrence network analysis

We employed the indicator species approach to investigate the future dynamics of vegetation change in the study area and evaluated the likelihood of abrupt changes occurring in the future. Indicator species often exhibit sensitivity to environmental stressors, and changes in them can readily reflect the biotic or abiotic state of the environment, reveal evidence for the impacts of environmental change, or indicate the diversity of other species, taxa, or communities within an given area^63^. To select an appropriate indicator taxon for reflecting vegetation change, the co-occurrence network analysis was performed. The co-occurrence network was established based on the pairwise Pearson’s correlation matrix, which was calculated using the R package *psych*^64^. Links with an absolute value of the correlation coefficient (*r*) greater than 0.2 that were statistically significant (*p* < 0.05) were included in the co-occurrence network.

Degree, betweenness, closeness, and eigenvector centrality metrics were calculated to select indicator taxon in the pollen assemblages. The degree of a node refers to the number of edges directly connected to it, reflecting the node’s direct influence or activity within the network. Betweenness measures a node’s ability to act as a ‘bridge’ in a network, thereby influencing the flow of information. Closeness is calculated as the inverse of the average shortest path length from one node to all other nodes in a network, indicating the node’s efficiency in interacting with others. Eigenvector is equivalent to the weighted sum of the centrality scores of its neighboring nodes, reflecting the node’s overall importance within the network. Indicator taxon was identified by high degree, betweenness, closeness, and eigenvector centrality. To minimize the bias introduced by pollen identification and avoid an overly complex network, 34 major terrestrial pollen taxa (with a mean percentage abundance ≥ 0.1%) were included in the construction of the co-occurrence network. Co-occurrence network analysis and the calculation of network metrics degree, betweenness, closeness, and eigenvector centrality) were conducted using R package *igraph*^65^.

### 3.6 Species distribution models

We used species distribution models to simulate the potential distribution of the indicator taxon *Tsuga* within HMs across past periods and under various future climate scenarios. This analysis comprised four main steps: preparing taxon occurrence data, selecting environmental variables, constructing and validating models, and generating predictions:

*i) Occurrence data collection.* The occurrence data of *Tsuga* in HMs were sourced from the Global Biodiversity Information Facility (GBIF, https://www.gbif.org/) and vegetation survey data collected by our research group (Fig. S7). In total, 475 modern distribution record were incorporated, with 248 of them originating from GBIF.
*ii) Bioclimatic variables and selection.* Bioclimate data were sourced from the World Climate Database (WorldClim, https://www.worldclim.org/) at a spatial resolution of 30 second, encompassing 19 bioclimate variables for both present and future periods. Data for future periods (2021–2040, 2041–2060, 2061–2080, and 2081–2100 CE) were derived from four Shared Socioeconomic Pathways (SSP126, SSP245, SSP370, SSP585) of five models (ACCESS-CM2, CMCC-ESM2, EC-Earth3-Veg, MRI-ESM2-0, UKESM1-0-LL) from the Couple Model Intercomparison Project Phase 6 (CIMP6). Bioclimate data for past periods (12.6–15.0, 15.1–17.3, 17.4–18.5, 18.6–19.6, 19.7–22.0 cal kyr BP) were derived from the CHELSA-TraCE21k dataset^40^. We used the bioclimate data for the present period (average for 1970–2000 CE), along with the occurrence data of *Tsuga*, to construct, calibrate, and validate species distribution model. To reduce the impact of overfitting due to collinearity among bioclimate variables, a threshold-based (Pearson’s pairwise correlation coefficient) pre-selection approach was used in this study to remove correlated variables. This analysis was conducted by the function *removeCollinearity* from R package *virtualSpecies*^66^. Finally, 7 bioclimatic variables with correlation coefficient were selected, including BIO2 (mean diurnal range), BIO3 (isothermality), BIO4 (temperature seasonality), BIO10 (mean temperature of warmest quarter), BIO12 (annual precipitation), BIO15 (precipitation seasonality), and BIO17 (precipitation of driest quarter) (Fig. S8).
*iii) Species distribution model building and validation.* We initially used eleven modelling algorithms from the R package *biomod2*^67^ to construct species distribution models for *Tsuga* in HMs. These algorithms included artificial neutral network (ANN), classification tree analysis (CTA), Flexible discriminant analysis (FDA), generalized additive model (GAM), generalized boosting model (GBM), generalized linear model (GLM), multiple adaptive regression splines (MARS), maximum entropy (MAXENT), random forest (RF), surface range envelop (SRE), and extreme gradient boosting training (XGBOOST). The occurrence data were randomly partitioned into an 80% sample for model calibration and the remaining 20% for model performance evaluation. The robustness of these models was assessed using threshold-dependent true skill statistics (TSS)^68^ and threshold-independent receiver operating characteristics (ROC) of area under the curve^69^. Based on the TSS and ROC metrics, we ultimately selected the RF model as it was the only model with both ROC and TSS values exceeding the threshold of 0.8 (Fig. S9), indicating good to excellent model performance.
*iv) Potential distribution predicting.* Potential distributions of *Tsuga* within the Hengduan Mountains for both past and future periods were predicted using the established species distribution model and the selected bioclimatic variables. The prediction values, which ranges between 0 and 1, were reclassified into four suitability categories: extreme suitability (> 0.8), high suitability (0.6–0.8), moderate suitability (0.4–0.6), low to no suitability (< 0.4).

The temporal patterns of changes in the proportions of extremely and highly suitable habitats for *Tsuga* across the past periods generally align with those revealed by the percentage abundance of *Tsuga* in the pollen record (Fig. S6), thereby providing additional validation for the model’s predictive accuracy. To identify trends in *Tsuga*’s distribution within HMs under different future scenarios by the end of this century, we calculated Theil-Sen estimator (also called Sen’s slope estimator) for each grid cell based on the simulations of the four future periods. Sen’s slope estimator is a non-parametric method for estimating the slope of a time series, defined as the median of the slopes of all possible pairs of data points^70^. This approach is robust to outliers and suitable for data that does not follow a normal distribution^71^. A higher absolute value of the estimated Sen’s slope estimator indicates a faster rate of change in *Tsuga*’s potential distribution. Positive and negative values of Sen’s slope reflect an increase or decrease, respectively, in the suitability for *Tsuga*’s distribution.

## CRediT Author Statement

M.L.: Conceptualization, Data curation, Formal analysis, Resources, Writing - Original Draft, Writing - Review & Editing.

P.L., Z.H., X.Z: Data curation, Writing - Review & Editing.

K.C., Y.W.: Investigation, Writing - Review & Editing.

X.L., E.Z., J.N.: Supervision, Resources, Writing - Review & Editing.

K.L.: Conceptualization, Investigation, Formal analysis, Resources, Writing - Original Draft, Writing - Review & Editing.

## Acknowledgement

This work is financially supported by the National Key Research and Development Program of China (2022YFF0801104) and the National Natural Science Foundation of China (42477484, 42472246). We appreciate the assistance from “大理白族自治州洱海管理局”.

## Supplementary Materials

**Figure S1.**
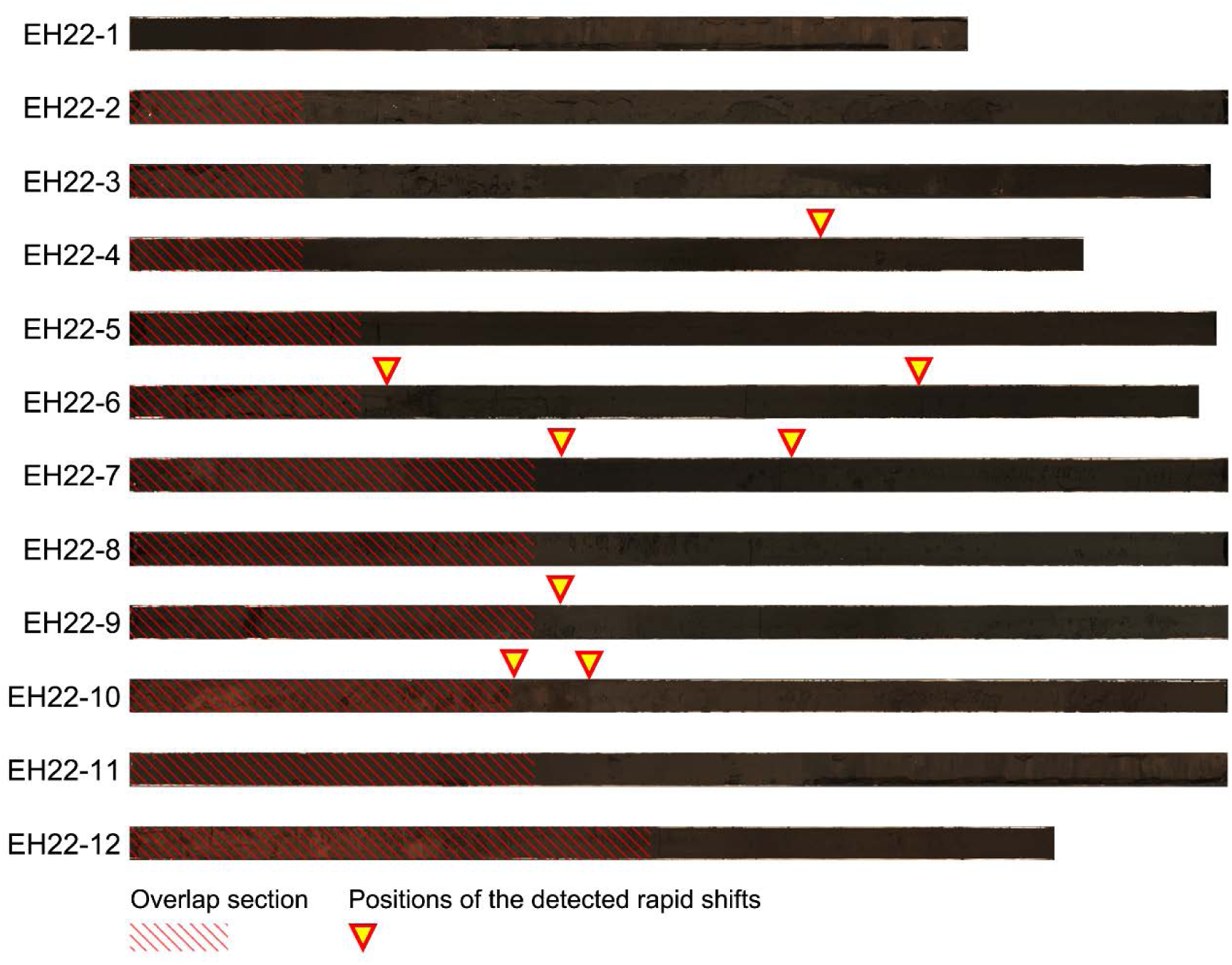
Photographs showing vertical profiles of the sediment core EH22. Inverted triangles indicate make the positions of eight rapid vegetation shifts (detected via pollen analysis; see Figure 3).

**Figure S2.**
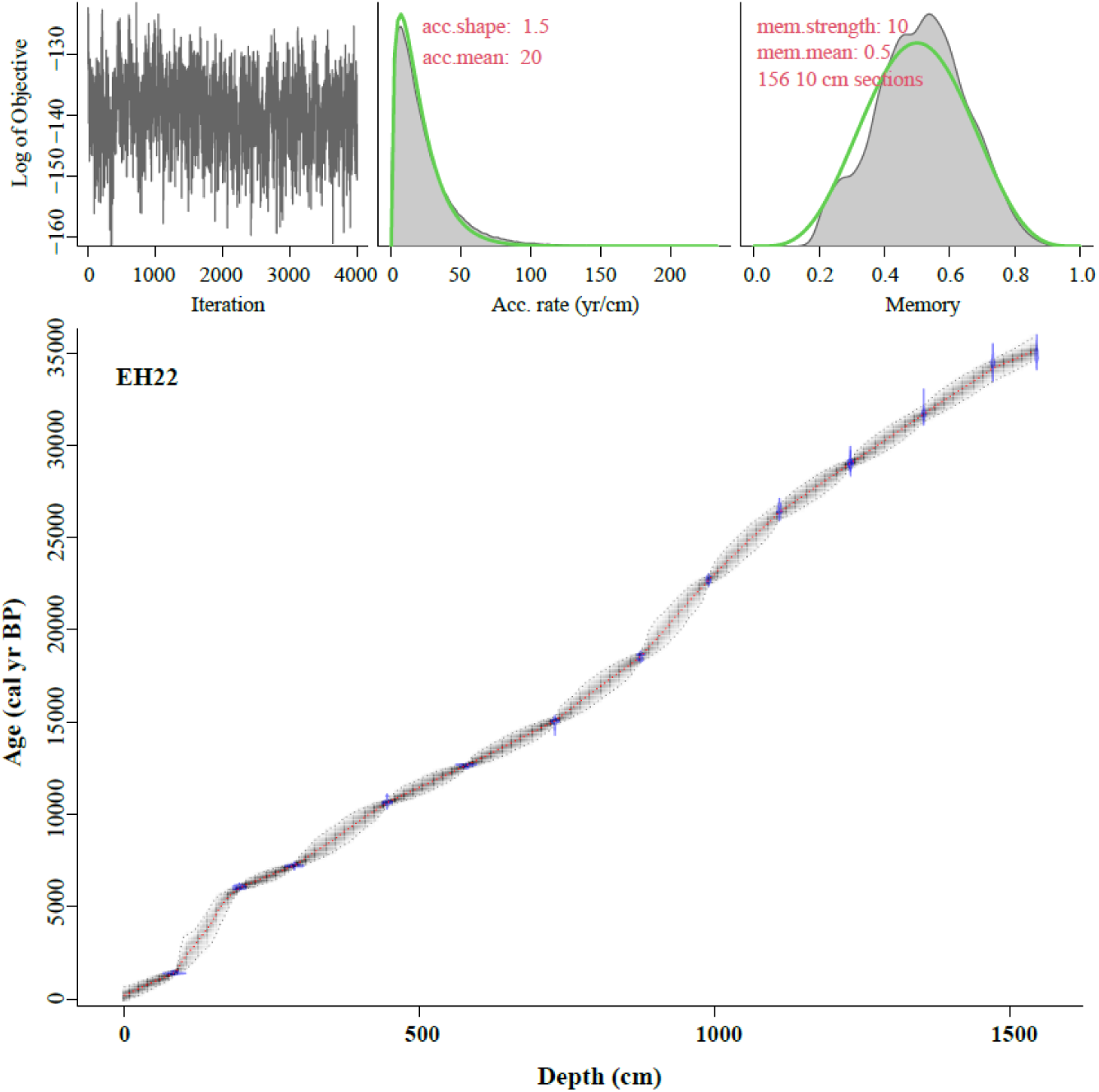
Age-depth model of core EH22. The model was established based on fourteen radiocarbon ages from different depths (see Table S1), with carbon reservoir effect of 1160 years.

**Figure S3.**
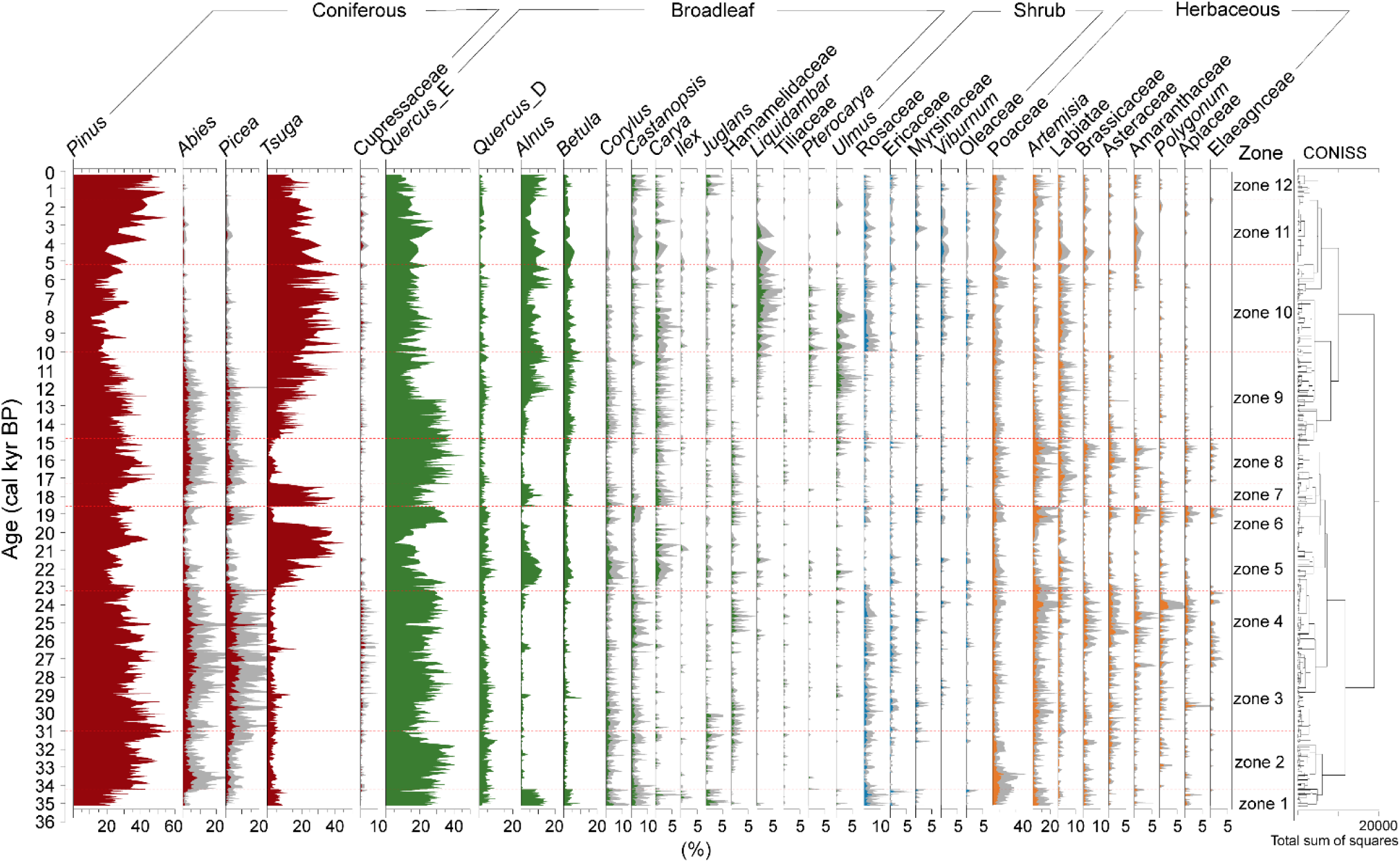
A percentage summary diagram of pollen taxa from the EH22 core over the past 35.2 cal kyr BP. Thirty-two major taxa (mean percentage higher than 0.1%) were displayed. The record was divided into twelve distinct zones using a stratigraphic cluster analysis based on the changes in the pollen assemblages.

**Figure S4.**
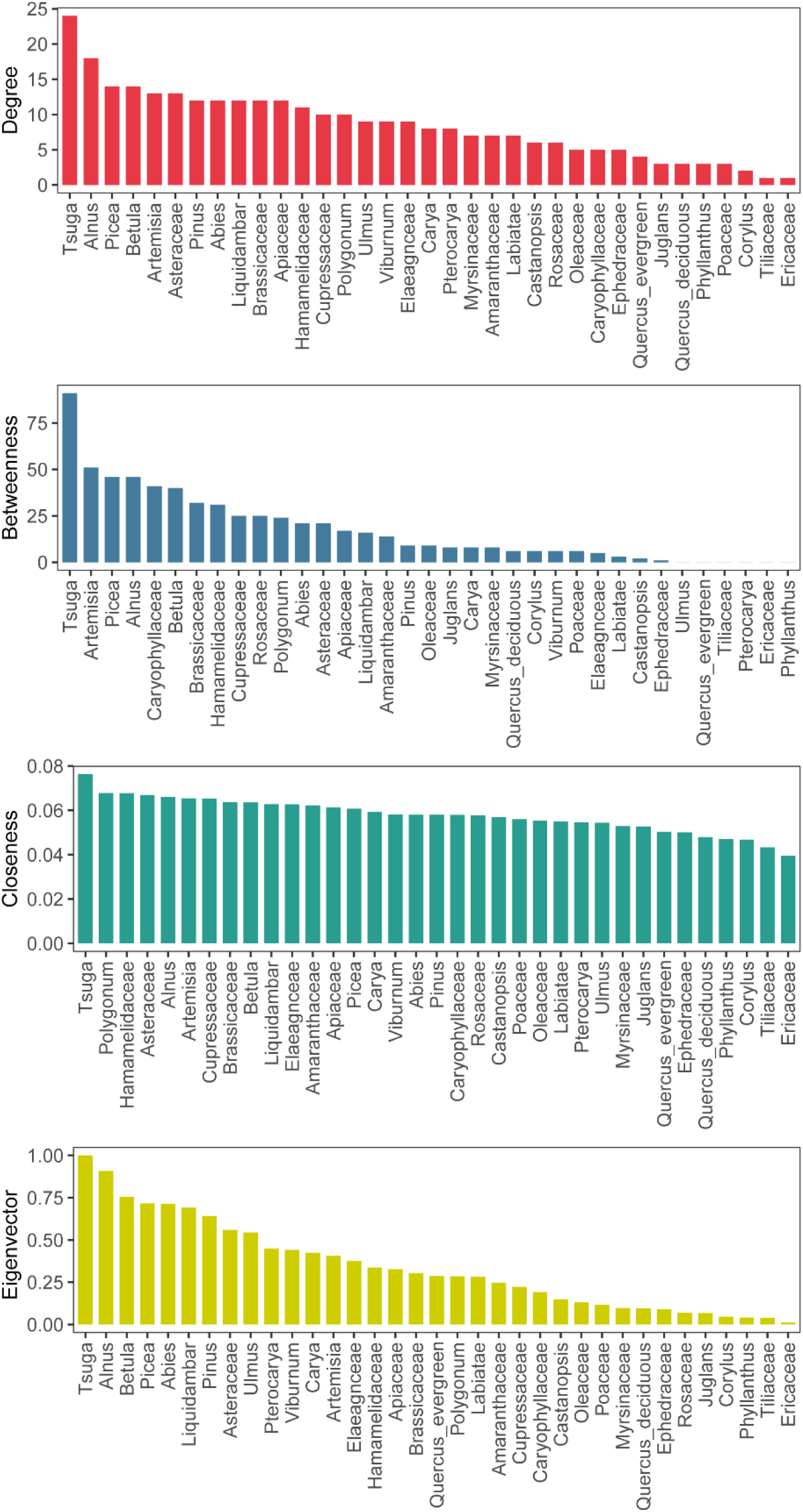
Node parameters (Degree, Betweenness, Closeness, Eigenvector) of the main pollen taxa estimated through network analysis.

**Figure S5.**
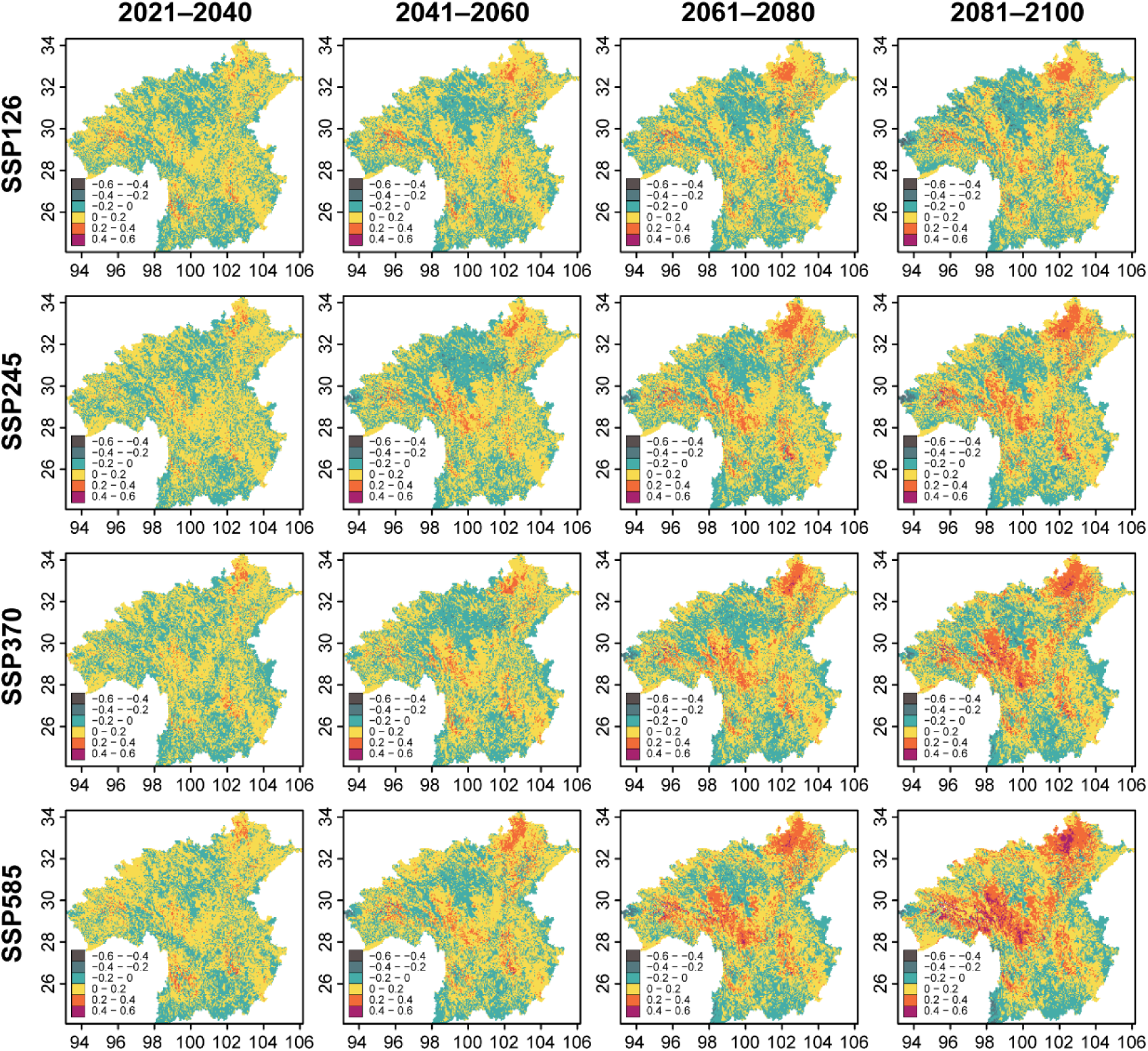
Projected changes in the habitat suitability for *Tsuga* spp. across the Hengduan Mountains (HMs) under four Shared Socioeconomic Pathway (SSP) scenarios (SSP126, SSP245, SSP370, SSP585) and four future periods (2021–2040, 2041–2060, 2061–2080, 2081–2100 CE). The baseline period is 1970–2000 CE.

**Figure S6.**
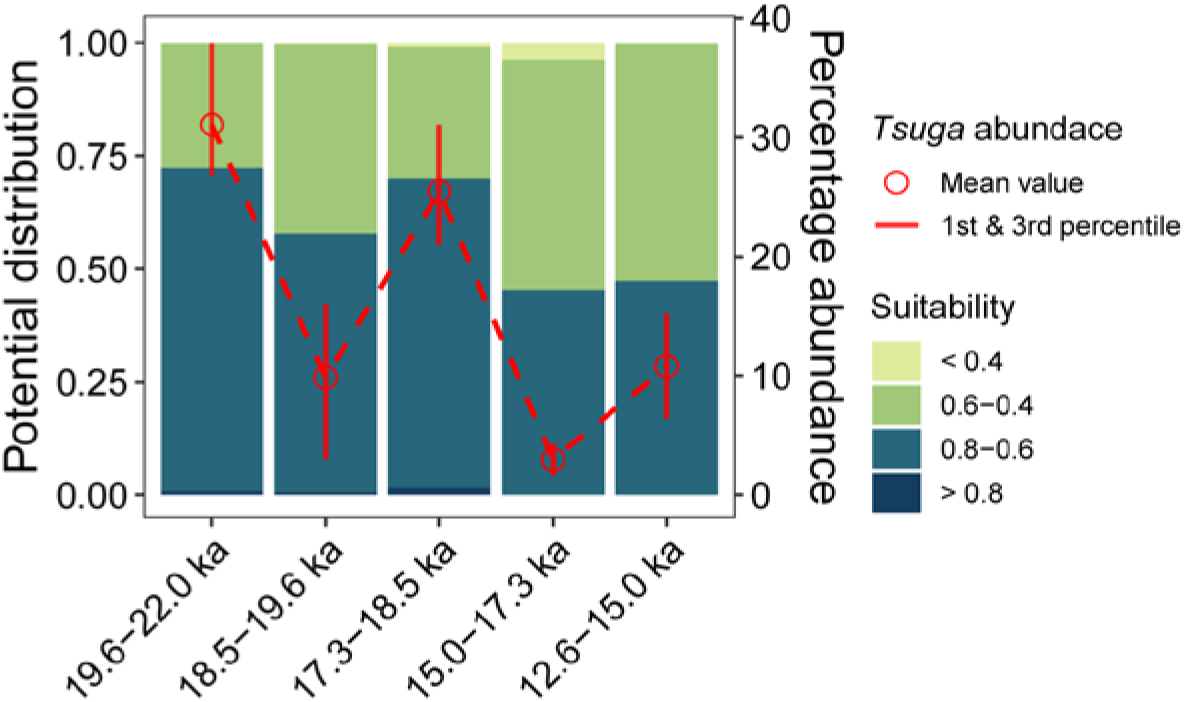
Projected proportions of *Tsuga* habitat suitability levels in the Erhai region (25.42–26.42°N, 99.88–100.43°E) compared with *Tsuga* abundance from sediment core EH22 across five historical periods.

**Figure S7.**
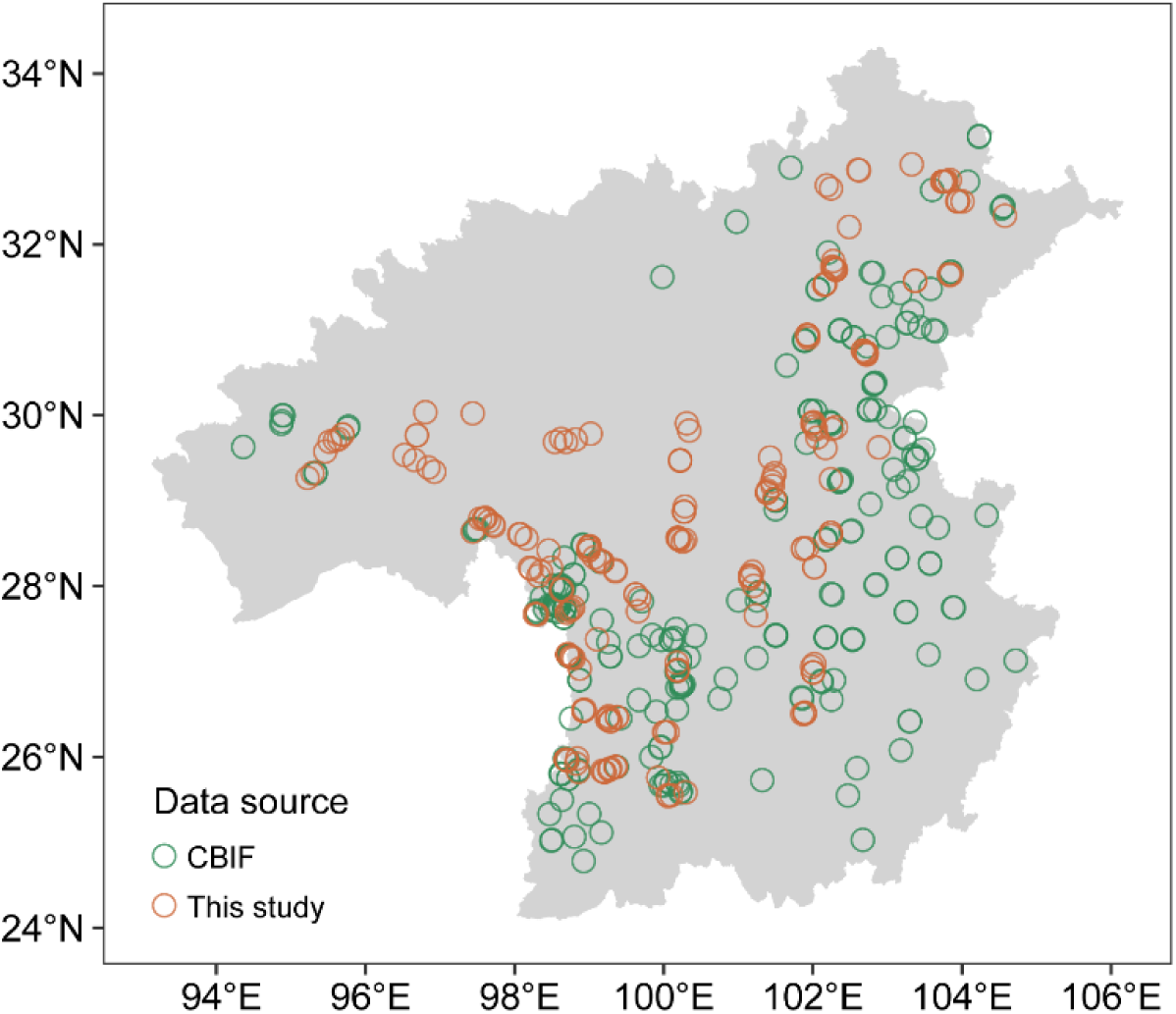
Data source and spatial distribution of the presence/absence information of *Tsuga* spp. in the Hengduan Mountains. The grey-shaded area represents the geographical extent of the mountainous region. CBIF: Global Biodiversity Information Facility (https://www.gbif.org/)

**Figure S8.**
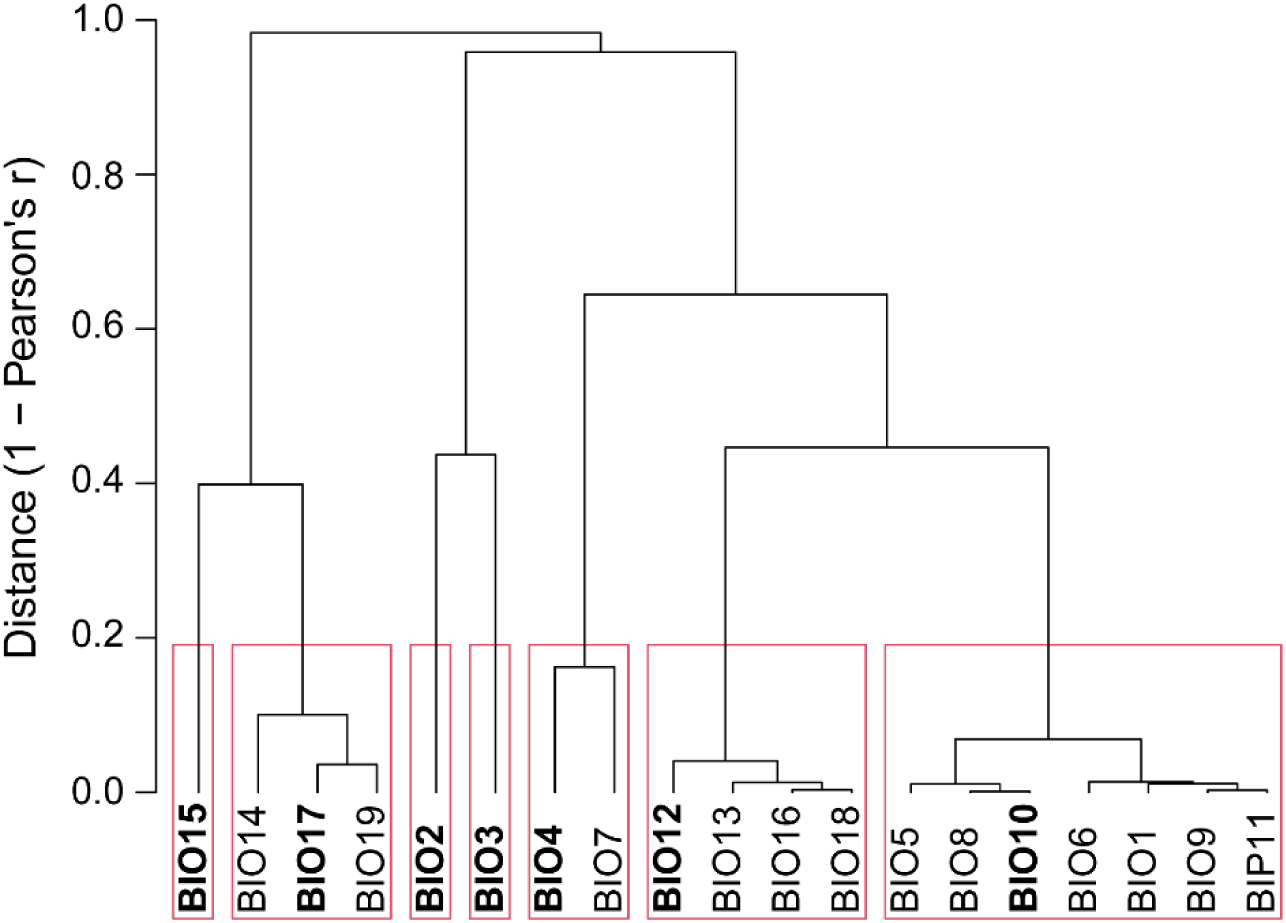
Cluster analysis of the timeseries for nineteen bioclimatic variables. Red rectangular boxes indicate groups of intercorrelated bioclimatic variables at a cutoff value of 0.7. Variables marked in bold represent the selected variables used in the species distribution model construction for *Tsuga*. Bioclimatic variables including: annual mean temperature (BIO01), mean diurnal range (BIO02), isothermality (BIO03), temperature seasonality (BIO04), max temperature of warmest month (BIO05), min temperature of coldest month (BIO06), temperature annual range (BIO07), mean temperature of wettest quarter (BIO08), mean temperature of driest quarter (BIO09), mean temperature of warmest quarter (BIO10), mean temperature of coldest quarter (BIO11), annual precipitation (BIO12), precipitation of wettest month (BIO13), precipitation of driest month (BIO14), precipitation seasonality (BIO15), precipitation of wettest quarter (BIO16), precipitation of driest quarter (BIO17), precipitation of warmest quarter (BIO18), precipitation of coldest quarter (BIO19).

**Figure S9.**
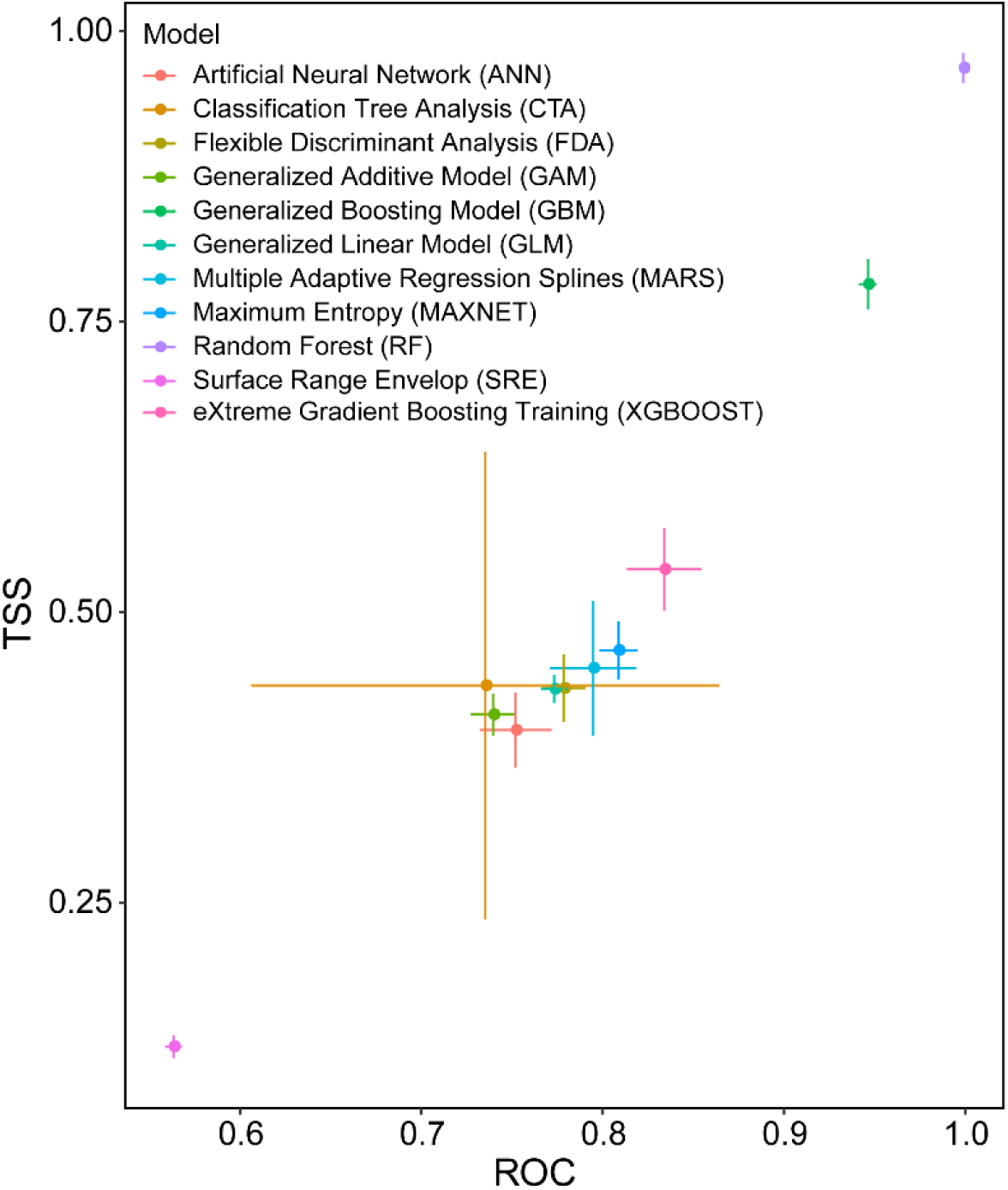
Mean evaluation scores (with their standard deviations) of receiver operating characteristics (ROC) and true skill statistics (TSS) for eleven species distribution models.

**Table S1.** The fourteen AMS radiocarbon dates of EH22 sediment core.

| Lab.ID<br>(Beta-) | Depth<br>(cm) | Materials | <sup>14</sup>C age<br>(yr BP) | Error<br>(± yr) | IRMS δ<sup>13</sup>C<br>(‰) | Calibrated <sup>14</sup>C age<br>(cal yr BP, 2σ) |
| --- | --- | --- | --- | --- | --- | --- |
| 639881 | 86 | bulk sediment | 2,520 | 30 | -24.6 | 2,493–2,600 |
| 639893 | 196 | bulk sediment | 6,260 | 30 | -28.8 | 7,156–7,263 |
| 639882 | 289 | bulk sediment | 7,310 | 30 | -28.3 | 8,030–8,177 |
| 639883 | 446 | bulk sediment | 10,420 | 40 | -27.0 | 12,097–12,491 |
| 639884 | 580 | bulk sediment | 11,650 | 40 | -25.8 | 13,438–13,597 |
| 639885 | 730 | bulk sediment | 13,620 | 40 | -23.8 | 16,301–16,606 |
| 639886 | 875 | bulk sediment | 16,290 | 50 | -24.3 | 19,526–19,846 |
| 639887 | 990 | bulk sediment | 19,800 | 60 | -23.8 | 23,728–23,982 |
| 639888 | 1110 | bulk sediment | 23,200 | 90 | -23.6 | 27,279–27,681 |
| 639889 | 1230 | plant remains | 24,730 | 110 | -9.5 | 28,751–29,175 |
| 642374 | 1230 | bulk sediment | 25,890 | 110 | -23.3 | 29,975–30,330 |
| 639890 | 1354 | bulk sediment | 28,790 | 150 | -23.6 | 32,741–33,775 |
| 639891 | 1471 | bulk sediment | 31,060 | 210 | -23.8 | 34,850–36,008 |
| 639892 | 1545 | bulk sediment | 31,480 | 190 | -23.9 | 35,404–36,219 |

